# Human embryo implantation involves Syncytin-2/MFSD2A-mediated heterokaryon formation with maternal endometrium

**DOI:** 10.64898/2025.12.22.695952

**Authors:** Tomas E. J. C. Noordzij, Martina Celotti, Ruben van Esch, Lisa Sackmann, Adriana Martìnez-Silgado, Franka de Jong, Hiromune Eto, Harry Begthel, Jeroen Korving, Theresa M. Sommer, Gaby S. Steba, Nicolas Rivron, Esther B. Baart, Johan H. van Es, Hans Clevers, Katharina F. Sonnen

## Abstract

Human embryo implantation involves attachment of the blastocyst to the endometrial epithelium to subsequently gain access to the underlying stromal compartment. The blastocyst is believed to cross the epithelium either by migration through, or upon apoptosis of, the endometrial epithelial cell layer. Yet, how the blastocyst exactly traverses the endometrial epithelium remains unknown. Here, we describe an *in vitro* implantation model of human blastoids and hormonally matured endometrial organoids amenable to high-resolution live imaging. We demonstrate that the initial step of implantation is mediated by the direct fusion of blastoid cells with endometrial epithelial cells. Blastoids express the fusion proteins Syncytin-1 and -2, while the endometrial epithelium mainly expresses the fusion co-receptor for Syncytin-2, called MFSD2A. CRISPR-induced loss of *MFSD2A* in the endometrial epithelium prevents blastoids from attaching and abolishes fusion. Together, these findings support a model in which fetal-maternal cell fusion constitutes the critical initiating mechanism of human embryo implantation, with endometrial *MFSD2A* playing an indispensable role in this process.

---

The early stages of human embryo implantation are critical for the successful establishment of pregnancy. Compared to most mammalian species, human fertility is relatively low, with approximately 60% of conception attempts failing, often for reasons that remain poorly understood. A key factor in achieving a successful early pregnancy is the proper attachment and invasion of the blastocyst into the endometrium, which happens between days 5 and 7 post-fertilization. During this period, the blastocyst enters the uterine cavity and aligns its polar trophectoderm towards the endometrium. Here, trophoblast cells within the polar trophectoderm differentiate and fuse to generate the multinucleated syncytiotrophoblast (STB)^2^. This cell fusion is mediated by the upregulation of the fusion proteins Syncytin-1 and Syncytin-2, which interact with their respective receptors, ASCT2 (*SLC1A5*) and MFSD2A^3–7^. The formation of the STB marks the initiation of implantation, as it coincides with breaching of the endometrial epithelium and subsequent invasion into the underlying stromal compartment - a crucial step in anchoring the embryo and establishing a functional placental interface. Two main mechanisms have been proposed to explain how the STB breaches the endometrial epithelium. One would involve transepithelial penetration, in which epithelial cells are displaced and engulfed^8,9^. The alternative hypothesis posits that the STB induces apoptosis of epithelial cells, thereby creating a gateway for entry into the underlying stroma^10^. However, current knowledge is largely derived from static observations, as methods enabling real-time analysis of human embryo implantation have not been available.

To investigate the early events of human embryo implantation, we sought to develop a co-culture system amenable to high-resolution imaging that could recapitulate the interaction between the blastocyst and endometrial epithelium (Fig.1a). To be able to distinguish the embryonic and endometrial components during imaging, we developed reporter lines with non-overlapping fluorescent reporters (Fig.1a). We established genetically engineered human endometrial organoids expressing Lifeact-mScarlet^11^ (Fig. 1b). To optimize imaging resolution and ensure an apical-out orientation, the endometrial organoids were transferred to a 2D culture format by placing the cells onto a thin coat (∼30µm) of hydrogel consisting of a mixture of collagen I and Matrigel® in a glass-bottom imaging chamber (Fig. 1c-d). This hydrogel mimics the collagen composition and stiffness of the *in vivo* stromal compartment^12,13^.

**Figure 1.**
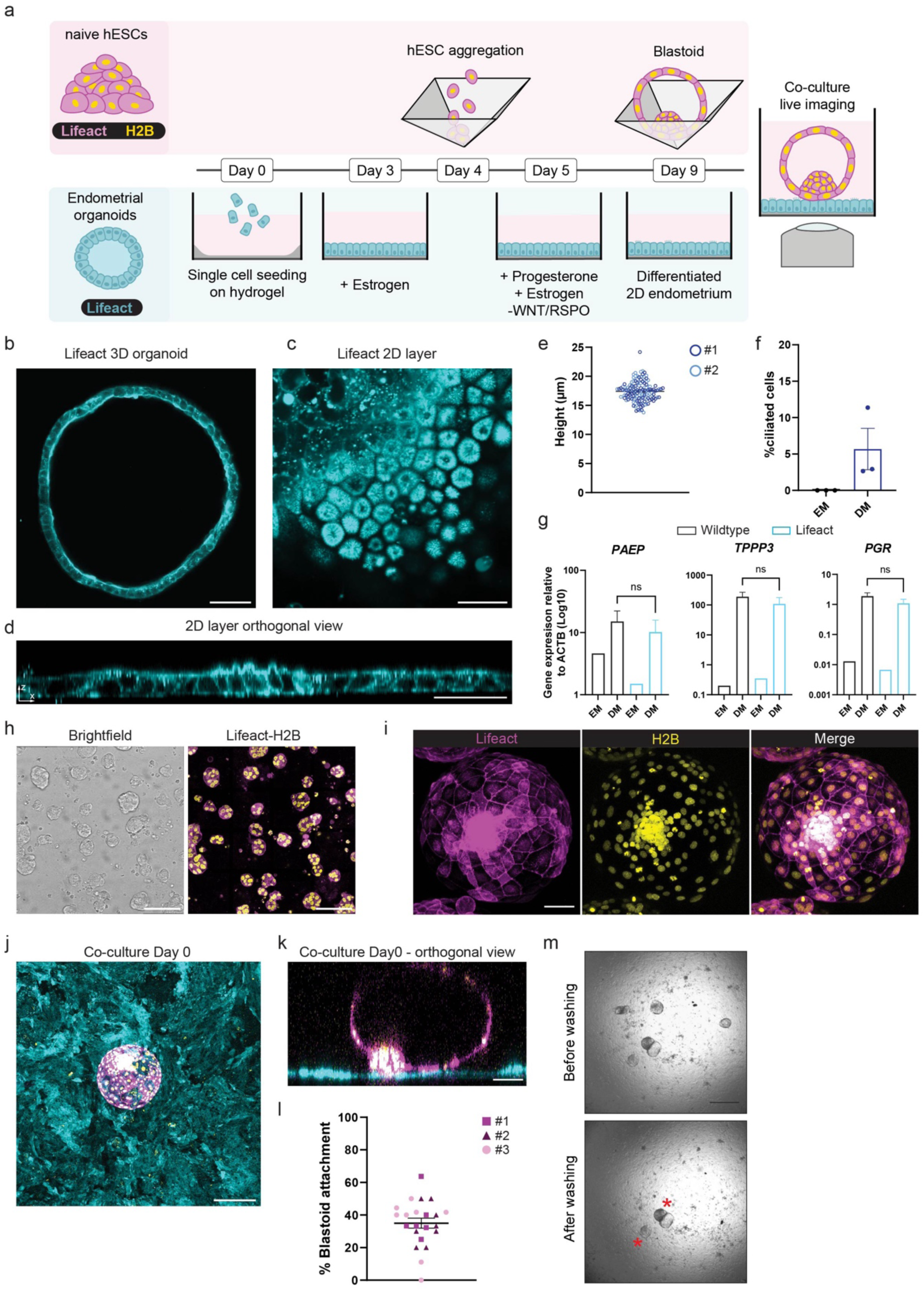
*In vitro* implantation setup amenable for high resolution imaging. **(a)** Schematic of the implantation setup amenable for live imaging. Top: generation of blastoids from Lifeact-GFP (magenta) and H2B-iRFP (yellow) expressing naïve human embryonic stem cells (hESCs). Bottom: generation of a 2D monolayer and differentiation of Lifeact-mScarlet (cyan) expressing endometrial organoids. **(b)** Representative image of a 3D Lifeact-mScarlet (cyan) expressing endometrial organoid. Scale bar, 50 µm. **(c)** Top view of a 2D Lifeact-mScarlet (cyan) expressing endometrial epithelial layer. Scale bar, 20 µm. **(d)** Orthogonal view of the 2D Lifeact-mScarlet (cyan) expressing endometrial layer. **(e)** Dot plot showing epithelial height from the basal to apical membrane (µm). Each dot represents a single cell; data represent mean ± s.e.m. (n = 2). **(f)** Quantification of ciliated cells (%). Comparison between expansion medium (EM) and differentiation medium (DM). Data represent mean ± s.e.m. (n = 3). **(g)** Relative expression of endometrial marker genes measured by RT-qPCR: *PAEP* (secretory cells), *TPPP3* (ciliated cells), and *PGR* (hormone-responsive gene). Comparison between Wildtype and Lifeact-mScarlet organoids cultured in EM or DM. Data represent mean ± s.e.m.; statistical analysis by unpaired *t*-test, two tailed, ns= non-significant. (n = 3). **(h)** Bright-field image of naïve hESCs. Fluorescence image showing Lifeact-GFP (magenta) and H2B-iRFP (yellow) expression. Scale bar, 50 µm. **(i)** Fluorescence images of a Lifeact-GFP (magenta) and H2B-iRFP (yellow) expressing blastoid, and merged channels. Scale bar, 50 µm. **(j)** Top view of day 0 of co-culture showing a Lifeact-GFP (magenta) and H2B-iRFP (yellow) expressing blastoid on a 2D Lifeact-mScarlet (cyan) expressing endometrial layer. Scale bar, 200 µm. **(k)** Orthogonal view of j. Scale bar, 50 µm. **(l)** Dot plot quantifying blastoid attachment (%). Each color represents an independent experiment; data represent mean ± s.e.m. (n = 3). **(m)** Bright-field images of co-cultures after two days, before and after washing. Red asterisks indicate attached blastoids. Scale bar, 500 µm.

The 2D organoids were stimulated with estrogen and progesterone to induce the secretory phase of the menstrual cycle^14,15^ (Fig. 1a). Similar to *in vivo* secretory endometrial epithelium, we found that this induced cells to become elongated with a height of ∼17,5 µm^16,17^ (Fig. 1e). The visualization of actin within these 2D organoids revealed distinct morphological changes typical of hormonal differentiation: the remodeling of cell-cell junctions and the emergence of elongated cells with sporadic apical protrusions resembling pinopodes^18^ (Extended data Fig. 1d, e, Video 1). Moreover, gene expression analysis and immunostainings revealed the presence of ciliated and secretory cells upon hormone stimulation of endometrial cells, essential for recreating the appropriate environment for implantation (Extended data Fig. 1a-c). The differentiation towards secretory endometrium yielded ∼6% of multi-ciliated cells in 2D endometrial organoids, similar to what has been found in receptive endometrium *in vivo*^19^ (Fig. 1f, Extended Fig. 1c). Importantly, Lifeact-mScarlet expression did not appear to affect differentiation, nor did we observe any morphological differences compared to the original 2D endometrial organoid line (Extended data Fig. 1f).

Separately, we generated human embryonic stem cells (hESCs) expressing Lifeact-GFP and H2B-iRFP, which were used to create blastoids to model the blastocyst as described previously^20,21^. Lifeact-GFP visualized cell borders and finer structures like apical microvilli, while H2B-iRFP marked nuclei (Fig. 1h, i, Extended data Fig. 2a). Expression of Lifeact-GFP and H2B-iRFP in blastoids did not appear to affect their development and morphology (Extended data Fig. 2a-d). The blastoids were cultured on the 2D monolayer of matured endometrial organoid cells. The three different fluorescent proteins allowed us to distinguish the blastoids from the endometrial cells (Fig. 1j, k). As blastocysts are believed to require 2-3 days to implant *in vivo*^22^, we co-cultured blastoids on the differentiated 2D endometrium for 72 hours. We then gently washed the monolayer with medium and counted the number of attached blastoids. We thus noted that between 20 - 40% of blastoids remained firmly attached (Fig. 1l, m), similar to what has been reported by other *in vitro* implantation experiments^20,23^.

Surprisingly, when examining the attached blastoids, we consistently observed the presence of large, multinucleated cells co-expressing Lifeact-mScarlet, Lifeact-GFP and H2B-iRFP (Fig. 2a, b). After 72 hours of co-culture, all attached blastoids contained such triple-positive, multinucleated cells. Given that (1) these cells contained Lifeact originating from both the blastoids and the endometrial cells and (2) that blastocysts are known to contain cells with fusogenic capacity, this suggested that blastoid cells may have fused with endometrial cells.

**Figure 2.**
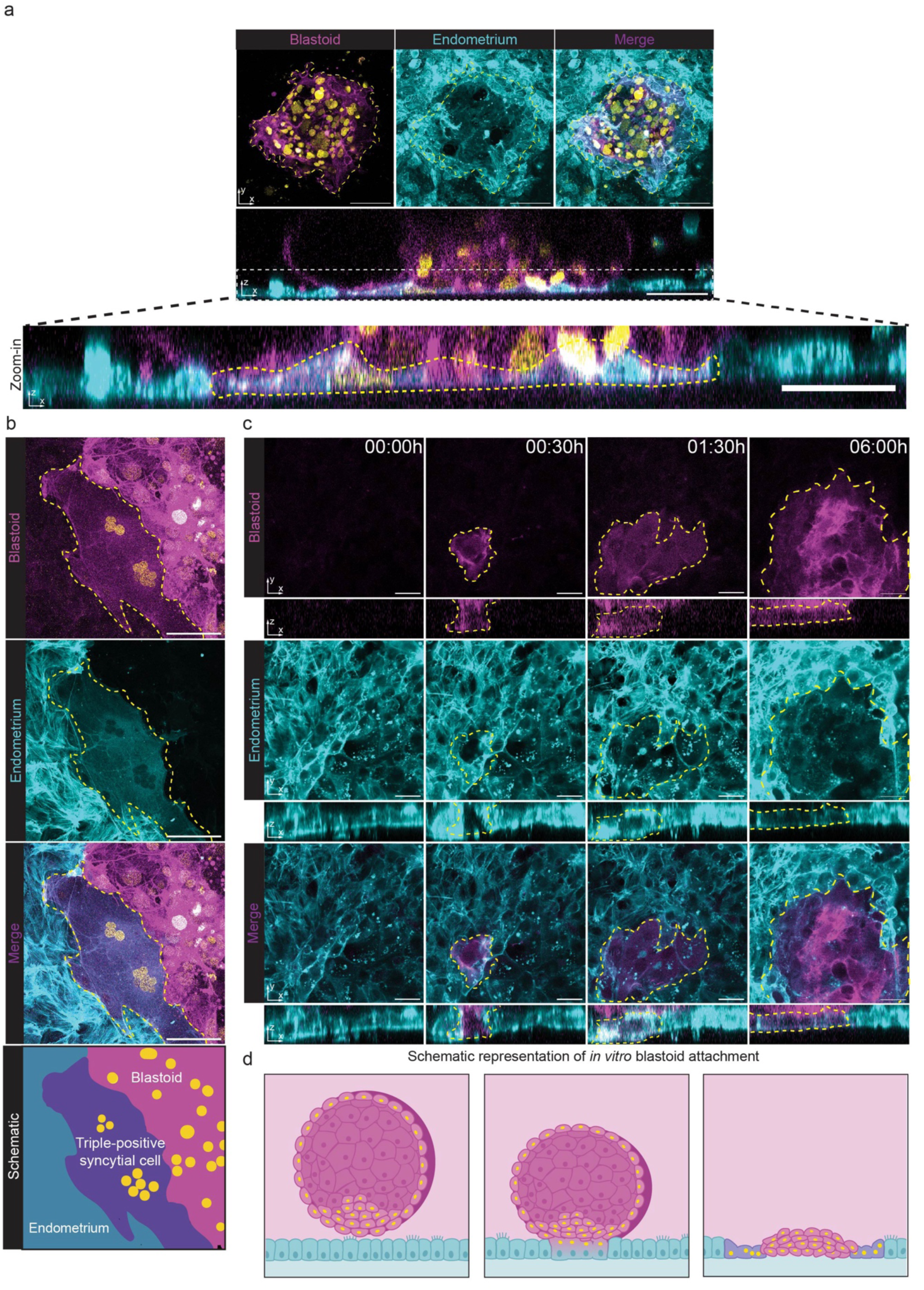
Blastoids fuse with the endometrial epithelium. **(a)** Representative images of the co-culture at day 2 showing fusion between the endometrium and blastoid. Top: top-view image showing the attached blastoid, surrounded by a giant cell positive for Lifeact-mScarlet (cyan), Lifeact-GFP (magenta) and H2B-iRFP (yellow). Bottom: orthogonal view of the same region, with a zoom-in (white box) highlighting the giant fused cell (surrounded by a yellow-dotted line). Scale bar, 50 µm. **(b)** Image representing a triple positive cell at the interface of the endometrium and blastoid (circled in yellow). Top panel: blastoid (magenta and yellow). Second panel: endometrium (cyan). Third panel: merged image showing fusion (purple), blastoid (magenta and yellow), and endometrium (cyan). Scale bar, 50 µm. On bottom, schematic illustration of the fusion (purple) between the endometrium (cyan) and blastoid (magenta and yellow) **(c)** Live imaging (top and orthogonal views) of blastoid-endometrial fusion. Top panels: blastoid (magenta). Middle panels: endometrium (cyan). Bottom panels: merged images showing the fused cells (circled in yellow). Scale bar, 20 µm. each image represents a z-plane of 1µm. **(d)** Schematic representation of *in vitro* blastoid attachment and fusion with the endometrial epithelium.

By comparing triple-positive cells in attached blastoids, we observed two morphologies: In ∼30 % of the attached blastoids, the triple-positive cells were located directly underneath the blastoid (group #1), while in the remaining ∼70 %, the triple-positive cells were located at the periphery of the interaction site between blastoid and endometrial monolayer, covering a larger area of the hydrogel (group #2) (Extended Fig. 3a-c). To understand the dynamics of this fusion event and the relationship between the two morphologies, we performed over-night live imaging with a time resolution of 30 minutes to capture the moment of blastoid attachment and the subsequent events. Attachment appeared to be initiated by an acute fusion event between the blastoid and endometrium (Fig. 2c, d, Video 2). First, the blastoid fused with one endometrial cell, which was rapidly followed by fusion with multiple neighboring endometrial cells. This resulted in the formation of a syncytial cell that quickly increased in size over the following six hours, after which further expansion slowed down (Extended data Fig. 4a, b). After 21 hours of imaging, we observed that further blastoid cells penetrated through the syncytial cell layer without further fusion, resulting in triple-positive cells moving towards the periphery (Fig. 2d). After an extended co-culture of 8 days, the triple-positive cells were retained at the interface of the blastoid and the endometrium (Extended data Fig. 3d). It thus appeared that the first morphology (group #1) represented the early stage of blastoid attachment, while the second (group #2) represented the stage in which the blastoid had attached via fusion and started to spread further on the hydrogel. We confirmed the fusion event using endometrial organoids from an independent donor (Extended data Fig. 3e).

Notably, following fusion between blastoids and endometrium, nuclei started appearing that were originally negative for the blastoid iRFP histone mark (and therefore were of endometrial origin), yet became progressively positive for the mark (Extended data Fig. 5a, b, Video 3). This observation indicated that histones encoded in the blastoid nuclei integrated into endometrial nuclear chromatin and further supported the notion that endometrial and blastoid nuclei resided in the same cells. To confirm these observations, we generated blastoids stably expressing H2B-mNeon and endometrial organoids expressing H2B-iRFP (Extended data Fig. 5c). After 72 hours of co-culture, we observed clusters of nuclei that were positive for both H2B-mNeon and H2B-iRFP (Extended data Fig. 5d, e). F-actin staining confirmed that the double-positive nuclei resided within the same cell (Extended data Fig. 5f). Within these fused cells, variable fluorescence intensities were observed per nucleus. Nuclei with higher H2B-mNeon intensity were presumably blastoid-derived, while those with higher H2B-iRFP intensity were endometrium-derived (Extended data Fig. 5g). Of note, individual syncytia consistently contained multiple trophoblast and endometrial nuclei.

The core machinery driving STB formation in the blastocyst consists of four proteins: the fusion ligands Syncytin-1 (*ERVW-1*) and Syncytin-2 (*ERVFRD-1*), and their respective receptors ASCT2 (*SLC1A5*) and MFSD2A *(MFSD2A)*^6,24^ (Fig. 3a). Immediately after formation of blastoids, in the absence of endometrium, Syncytin proteins were expressed at low levels, while higher expression levels were detected for the two receptors (Fig. 3b). After 2 days of expanded blastoid culture in the absence of endometrium, qPCR gene expression analysis revealed a significant upregulation of the ligands *ERVW-1* and *ERVFRD-1*, while the receptor *SLC1A5* expression remained stable and *MFSD2A* showed a modest increase (Fig. 3b). Notably, the expression of the STB marker genes *OVOL1*, *GCM1* and *GCB* were increased after two days, and we confirmed the presence of multinucleated cells within the blastoids (Fig. 3c, d). These findings indicate that under the given culture conditions, blastoid cells were capable of maturing and differentiating into STB cells.

**Figure 3.**
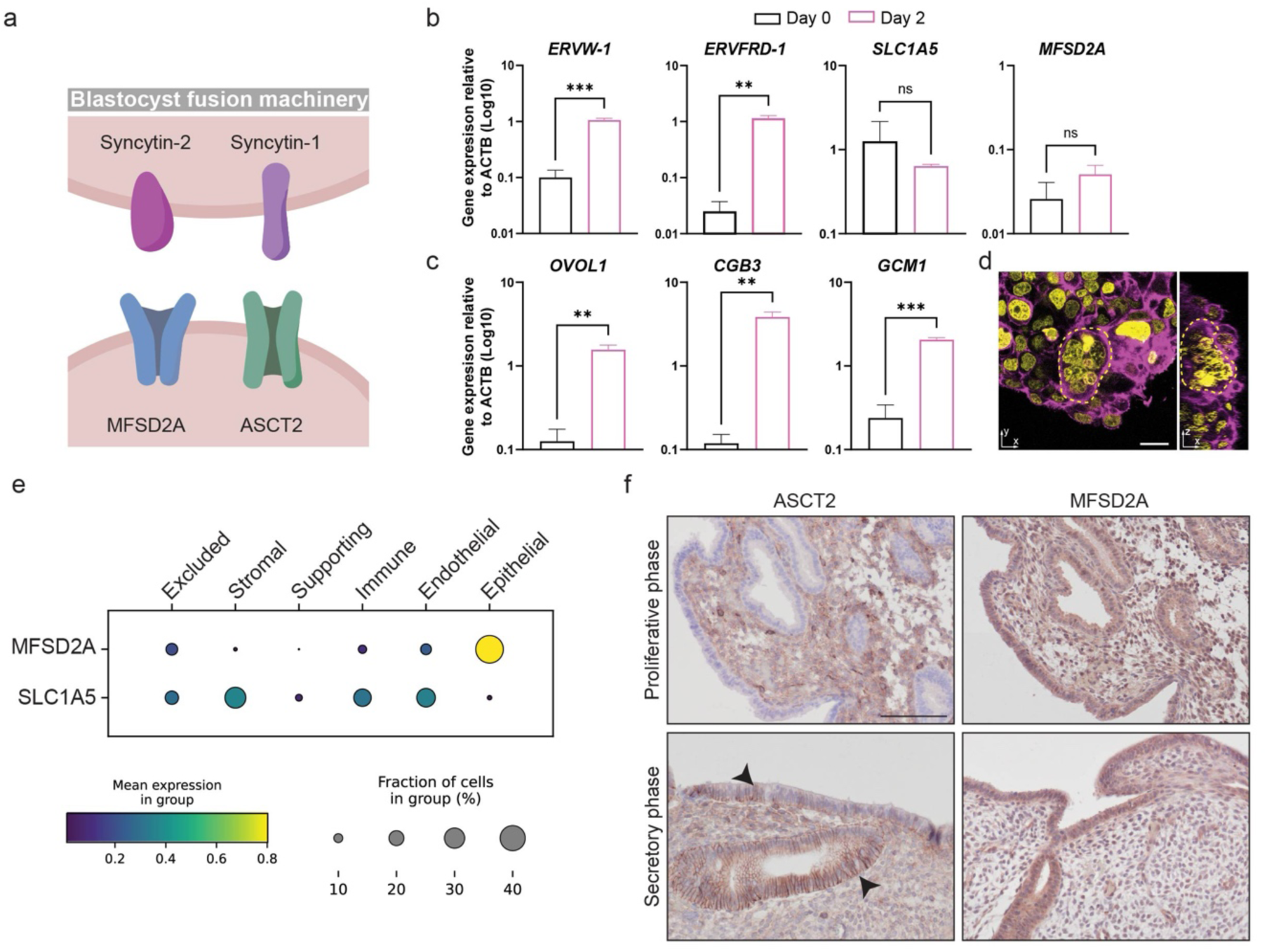
Fusion proteins are present in blastoids and *in vivo* endometrial tissue. **(a)** Schematic representation of the fusion machinery in the blastocyst, highlighting fusion ligands (Syncytin-2 (*ERVFRD-1*) and Syncytin-1 (*ERVW-1*)) and their receptors (MFSD2A and ASCT2 (*SLC1A5*)). **(b)** Relative expression of fusion genes (*ERVW-1*, *ERVFRD-1*, *SLC1A5* and *MFSD2A*) in blastoids at day 0 and day 2 after formation, measured by RT-qPCR. Data represent mean ± s.e.m.; statistical analysis by unpaired *t*-test, two tailed, ns= non-significant, *** (p-value= 0.0004), ** (p-value=0.0014) (n = 3). **(c)** Relative expression of syncytiotrophoblast marker genes (*OVOL1*, *CGB3* and *GCM1*) in blastoids at day 0 and day 2 after formation, measured by RT-qPCR. Data represent mean ± s.e.m.; statistical analysis by unpaired *t*-test two tailed, ** (p-value< 0.005), **** (p-value<0.0005) (n = 3). **(d)** Fluorescence image of a Lifeact-GFP (magenta) and H2B-iRFP (yellow) blastoid at the polar side, two days after formation, showing a syncytiotrophoblast (circled in yellow). Scale bar, 20 µm. **(e)** Dot plot showing *MFSD2A* and *SLC1A5* expression in in vivo endometrium. Bottom right: viridis color scale indicating mean expression per cell group. Bottom left: proportion of cells expressing each gene (percentage). Data analyzed from the dataset by García-Alonso *et al.* **(f)** Immunohistochemistry of endometrial tissue for ASCT2 and MFSD2A, comparing proliferative and secretory phases. Arrows indicate regions with SLC1A5 expression. Scale bar, 50 µm.

The upregulation of STB markers coincided with the time window during which blastoid–endometrial fusion was observed in our previous experiments. Since STB formation and epithelial fusion appeared to occur concurrently, it was plausible that the Syncytin-1:ASCT2 and/or the Syncytin-2:MFSD2A interactions could also mediate blastoid-endometrial fusion. To explore this, we examined the expression of these receptors *in vivo* using the single-cell RNA-seq dataset generated by Garcia-Alonso et *al*.^25^. This analysis revealed that *MFSD2A* is strongly expressed in endometrium epithelial cells, specifically in luminal and SOX9+ cells, whereas *SLC1A5* is predominantly localized to the stromal compartment of the endometrium (Fig. 3e). Immunohistochemical staining of primary human endometrial tissue confirmed these findings: MFSD2A was consistently expressed in epithelial cells throughout the proliferative and secretory phases, while ASCT2 was confined to stromal cells during the proliferative phase and to glandular epithelium during the secretory phase (Fig. 3f). Encouragingly, *MFSD2A* expression occurred in the 2D endometrial epithelial system and was upregulated upon maturation towards the secretory phase (Fig.4a).

To functionally assess the requirement of MFSD2A for blastoid-endometrial fusion, we generated clonal endometrial organoid lines lacking functional MFSD2A by introducing a premature stop codon using CRISPR base-editing (Extended Data Fig. 6a). Successful mutation was confirmed by Sanger sequencing (Fig. 4b). MFSD2A^KO^ organoids displayed similar morphology and long-term growth to their wildtype counterparts (Fig. 4c). Gene expression analysis further showed that MFSD2A^KO^ organoids retained normal differentiation capacity, displaying upregulation of lineage markers such as *PAEP* (secretory cells), *TPPP3* (ciliated cells), and *PGR* upon hormone-induced differentiation (Fig. 4d, Extended Data Fig. 6b). Next, we cultured blastoids for 72 hours on 2D endometrial epithelium derived from either wildtype or clonal MFSD2A^KO^ organoids. Strikingly, whereas blastoids readily attached to the wildtype endometrium, no stable adhesion was observed in the MFSD2A^KO^ condition (Fig. 4e). Blastoids cultured on mutant endometrial 2D organoids did not remain adherent upon medium flushing, indicating a loss of stable interaction with the endometrium (Fig.4f). We concluded that fusion did not occur between blastoids and the MFSD2A^KO^ endometrial epithelium, and that fusion is a prerequisite for stable attachment. Together, these results identified MFSD2A as an essential epithelial receptor required for both attachment and fusion with blastoids (Fig. 4g).

**Figure 4.**
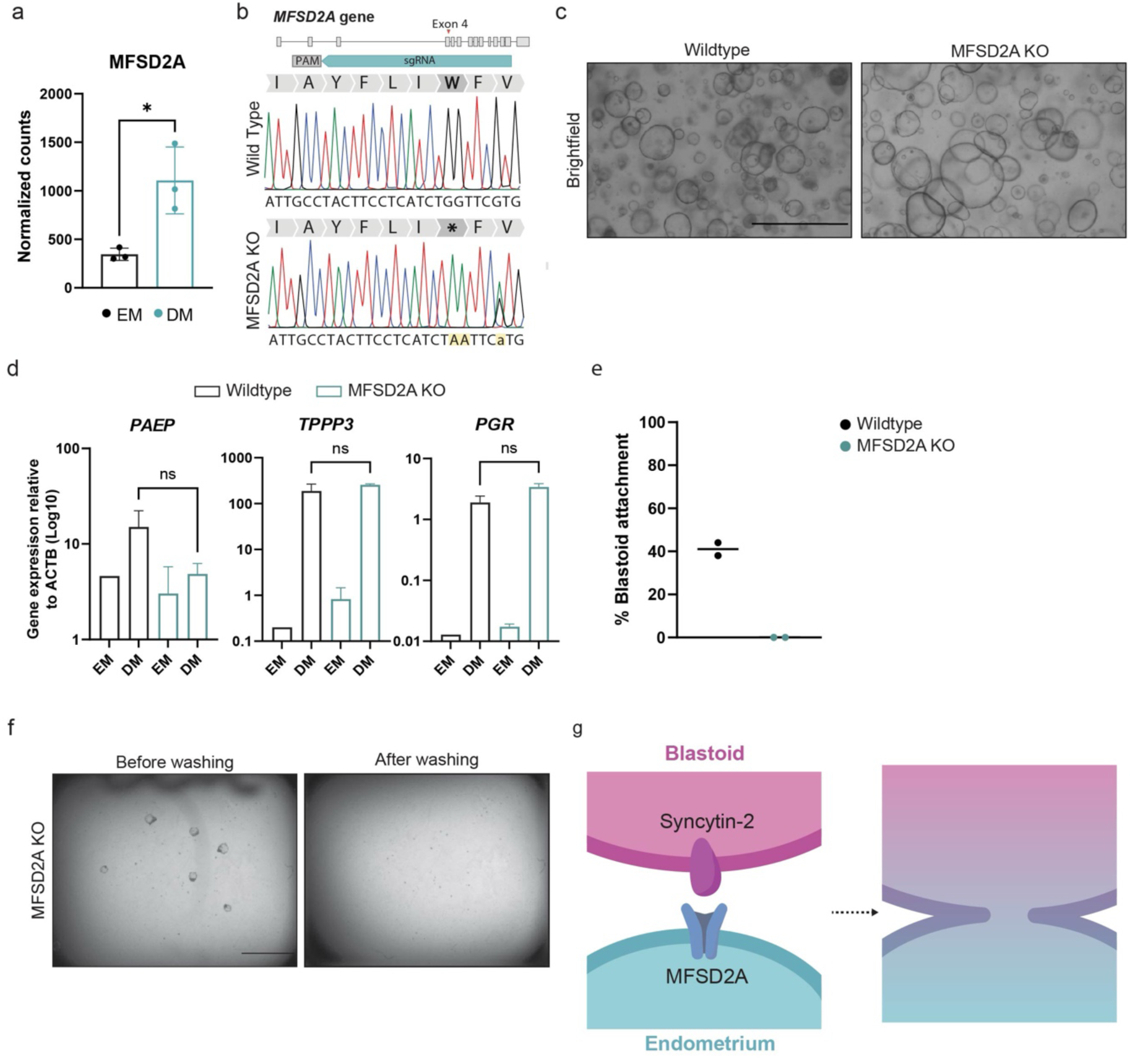
Fusion mediated by MFSD2A is essential for blastoid attachment to the endometrial epithelium. **(a)** Expression of *MFSD2A* (normalized counts) in endometrial organoids cultured in expansion medium (EM) or differentiation medium (DM). Data represent mean ± s.e.m.; statistical analysis by unpaired *t*-test two tailed, * (p-value= 0.0448) (n = 3, technical replicates). **(b)** Validation of *MFSD2A* knockout in endometrial organoids by Sanger sequencing, showing a tryptophan-to-stop codon mutation in exon 4 (W134*). **(c)** Bright-field images of Wildtype and *MFSD2A* knockout (KO) endometrial organoids. Scale bar, 1 mm. **(d)** Relative expression of endometrial marker genes measured by RT-qPCR: *PAEP* (secretory cell marker), *TPPP3* (ciliated cell marker), and *PGR* (hormone-responsive gene). Comparison between Wildtype and *MFSD2A* KO organoids cultured in EM or DM. Data represent mean ± s.e.m.; statistical analysis by unpaired *t*-test, two tailed, ns= non-significant (n = 3). **(e)** Blastoid attachment efficiency (%) in co-cultures with Wildtype (pink) or *MFSD2A* KO (green) endometrial organoids. Data represent mean ± s.e.m. Total of 55 blastoids analyzed for the knockout condition and 71 for wildtype. (n = 2). **(f)** Bright-field images of co-cultures with *MFSD2A* KO organoids before and after washing, showing attached blastoids. **(g)** Schematic model of the proposed fusion mechanism.

Taken together, by combining fluorescently labeled human blastoids and endometrial organoids in an *in vitro* implantation model amenable for high spatiotemporal-resolution live imaging, we have visualized an unexpected, highly dynamic interaction between human blastoids and the endometrial epithelium. The described fusion mechanism is distinct from earlier models that suggest that the endometrial epithelium undergoes apoptosis or passively retracts to permit embryo invasion. While these processes may coexist, our findings highlight cell fusion as a necessity for stable attachment, emphasizing the active role of the endometrium in early implantation. Of note, it has recently been shown that stromal cells can fuse with the STB *in vitro*, suggesting that multiple maternal cell types can contribute to the placenta^23^. Several observations support that fusion does occur *in vivo.* For instance, the primary syncytium contains differently sized nuclei: larger ones formed by endoreduplication, and smaller ones, potentially originating from the endometrial epithelium or stroma. Moreover, the concept of fusion of the blastocyst with the endometrial epithelium is not entirely new, as an early study in 1971 has reported that rabbit blastocysts may fuse with the endometrial epithelium^26^. Similar to what we have observed, fusion with the endometrial epithelium occurred in a localized fashion and did not spread along the entire epithelium. Interestingly, rabbits express only one Syncytin, called *Syncytin-Ory1*, orthologous to human Syncytin-1, which binds to the orthologous receptor ASCT2^27^. Our findings raise a plethora of questions. It will be important to understand whether fusion with endometrial cells indeed ensures proper anchoring to the uterus and whether the maternal cells contribute to the further invasion of the primary syncytium. An attractive hypothesis is that fusion with maternal cells could play a role in the evasion of the immune system. Finally, our findings imply that the endometrial epithelium is not just a barrier, but that it may play an active role during the initiation of pregnancy. Future studies of this atypical fusion may highlight why blastocysts fail to implant and could provide new therapeutic insights into human infertility.

## Acknowledgments

Research reported in this publication was supported by the European Research Council under an ERC starting grant agreement no. 850554 and by ZonMW under PSIDER grant number 10250022120003 (K.F.S, T.N. and L.S.) and the Hubrecht Institute. We thank the Hubrecht imaging center for the maintenance of equipment. We thank Bas van Rijn for providing tissue to establish endometrial organoid lines. We thank Daniel Krueger for providing the Lifeact-GFP, Lifeact-mScarlet and H2B-iRFP expression vectors. We thank Yasuhiro Takashima for providing us with the H9 cell line (WiCell Research Institute).

## Author contributions

T.N, M.C, H.C. and K.F.S. conceived the study; H.C. and K.F.S. supervised the project; J.v.E. and K.F.S. daily supervision; T.N and M.C. designed and performed the experiments; L.S., R.v.E., A.M.S. analysed data; H.E, F.d.J., H.B., J.K., E.B., G.S. helped with experiments; N.R. hosted T.N. to learn how to culture blastoids; T.N, M.C, H.C. and K.F.S. wrote the manuscript.

## Methods

### Ethical approval

The human embryonic stem cell line used in this study (H9W1) was previously derived and distributed with informed consent for research use, including genetic modification and differentiation. Fluorescent reporter lines were generated from the parental H9W1 line by random genomic integration of transgenes, in accordance with the approved protocol. This study does not involve the derivation of new human embryonic stem cell lines, the use of any newly obtained human biological samples, fetal or embryonic tissues, or donated human blastocysts. Furthermore, no *in utero* transfer of human-derived cells, tissues or embryo models into any species was performed. All research activities were conducted in full compliance with the ethical regulations of the Hubrecht Institute, Dutch legislation governing human embryonic stem cell research and the International Society for Stem Cell Research (ISSCR) Guidelines for Stem Cell Research and Clinical Translation (2021).

### Endometrial organoids culture

Human endometrial organoids used in this study were established from tissue biopsies as previously described and cryopreserved^15^. Established Organoids were maintained in expansion medium that contains: advanced DMEM-F12 medium (adDMEM/F12; Life Technologies), supplemented with 1 × GlutaMAX (11574466, ThermoFisher Scientific), 100 U/mL Penicillin–streptomycin (11548876, ThermoFisher Scientific), and 10 mM HEPES (11560496, ThermoFisher Scientific) [referred hereafter as AdDMEM+++]. AdDMEM+++, 2% final volume B27 Supplement (11530536, ThermoFisher Scientific), 1.25 mM N-acetylcysteine (A9165, Sigma-Aldrich), 10mM Nicotinamide (N0636, Sigma-Aldrich), 0.25% Noggin conditioned medium (U-Protein Express), 10% R-Spondin 1 conditioned medium (produced as described in Pleguezuelos-Manzano *et al.*^28^), 50 ng/ml EGF (AF-100-15, Peprotech), 100 ng/mL FGF10 (100-26, Peprotech), 1 µM A83-01 (2939, Tocris), 1 mM Prostaglandin E2 (2296, Tocris), Wnt3A surrogate (U-Protein Express), 10 mM ROCK inhibitor Y-27632 (M1817, Ab- mole) and 10 mg/mL Primocin (ant-pm-1, Invivogen). Organoids were maintained into a 37 °C, 5% CO2 incubator. The medium was changed every two to three days. Every 7 days organoids were dissociated through mechanical dissociation and passaged with a ratio 1:4.

### 2D culture and differentiation

Organoids were incubated in dispase for 30 min. After that, they were harvested and washed using AdDMEM+++. The organoid pellet was resuspended in 1 ml TrypLE Express (11568856, Thermo Fisher Scientific) and incubated at 37 °C for 5 min. This was followed by mechanical dissociation with P1000 until organoids were disrupted into single cells. These steps were repeated if required. Subsequently, 200,000 cells were seeded in 200 µL human endometrium expansion medium in an Ibidi well (8-well ibitreat high, Ibidi) on top of a gel mix of collagen (70%) (PureCol EZ Gel, Advanced Biomatrix) and Matrigel (30%). After 3 days, when cells had reached at least 70% confluency, differentiation was started: expansion medium was supplied with 10nM β-Estradiol (E1127, Sigma Aldrich) for 2 days to mimic the proliferative phase. After these 2 days, the medium was switched to differentiation medium for 5 days to mimic the secretory phase. Human endometrium differentiation medium consists of: AdDMEM+++, 2% final volume B27 Supplement (11530536, ThermoFisher Scientific), 1.25 mM N-acetylcysteine (A9165, Sigma-Aldrich), 10mM Nicotinamide (N0636, Sigma-Aldrich), 0.25% Noggin conditioned medium (U-Protein Express), 50 ng/ml EGF (AF-100-15, Peprotech), 100 ng/mL FGF10 (100-26, Peprotech), 1 µM A83-01 (2939, Tocris), 1mM Prostaglandin E2 (2296, Tocris), 1µM cyclic AMP (B7880, Sigma Aldrich), 200 ng/ml Progesterone (P0130, Sigma Aldrich), 10nM β-Estradiol (E1127, Sigma Aldrich) and 10 mg/mL Primocin (ant-pm-1, Invivogen).

### Generation of stable genetically modified organoids

To construct the Lifeact-mScarlet-Puro plasmid, the mScarlet coding sequence was amplified from Addgene plasmid #190658, with the Lifeact peptide sequence (MGVADLIKKFESISKEE) incorporated into the forward primer and inserted into the vector backbone (Addgene #240804) digested with XhoI and SmaI (Promega). The mT2TP transposase and H2B-iRFP-Blast plasmids were generated as previously described^29^. To establish the Lifeact organoid line, 5 µg of mT2TP transposase plasmid was co-transfected with 5 µg of Lifeact-mScarlet-Puro plasmid. Similarly, for introduction of H2B-iRFP into organoid lines, 5 µg of mT2TP transposase plasmid was co-transfected with 5 µg of H2B-iRFP-Blast plasmid. Organoids were dissociated into single cells as described above and electroporated in cuvettes (BTX) using NEPA electroporator (Nepa Gene). For selection of Lifeact-expressing organoids, cultures were treated with Puromycin (2 µg/mL; Invivogen). For selection of H2B-expressing organoids, Blasticidin (100 µg/mL; Invivogen) was used instead. Regarding the generation of Knockouts, the empty sgRNA plasmid backbone for SpCas9 was a kind gift from Keith Joung (BPK1520, Addgene plasmid #65777). MFSD2A W134* sgRNA sequence: 3’-CACGAACCAGATGAGGAAGT-5’. MFSD2A sgRNA was cloned as previously described^30^. Organoids were engineered using a C>T NGG base editor plasmid (SpCas9-CBE6b, Addgene plasmid #215280), and cells were electroporated as previously described^30^. One week after electroporation edited organoids were selected with addition of Hygromycin B Gold (100 µg/ml; Invivogen) to the medium. One week after selection, surviving organoids were picked and dissociated to single cells, and clonal lines were generated. To confirm the gene editing, DNA was extracted using Quick-DNA Microprep Kit (Zymo Research Corporation) and genotyped using gene-specific primer pairs. Forward 5’-TCCCAGTTCCCATCTGCCAT-3’, Reverse 5’-CACGATAGGCGGTGGCAGAA-3’ (sanger sequencing performed by Macrogen Europe BV). Successfully edited clonal lines were passed and cryopreserved.

### hESC culture

Human Naïve Embryonic Stem Cell lines H9W1, containing a Dox-inducible Gata6 expression vector, were kindly provided by Yasuhiro Takashima. In none of the experiments of this study, Gata6 expression was induced. Cells were cultured in PXGL medium, consisting of N2B27 basal medium including DMEM/F12 (50%, Gibco), neurobasal medium (50%, Gibco), N-2 supplement (Thermo Fisher Science, 17502048), B-27 supplement (Thermo Fisher Science, 17504044), 1x GlutaMAX supplement (Thermo Fisher Science, 35050-038), 1x non-essential amino acids (11140050, Gibco), 2-mercaptoethanol (100 µM, Thermo Fisher Science, 31350010), 1x Pen/Strep 5000 Units/ml, supplemented with PD0325901 (1 µM, MedChemExpress, HY-10254), XAV-939 (1 µM, MedChemExpress, HY-15147), Gö 6983 (2 µM, MedChemExpress, HY-13689) and human recombinant leukemia inhibitory factor (hLIF, 10 ng/ml, Stemcell, 78055). Cells were grown on irradiated mouse embryonic fibroblasts (MEFs, made in house) with a coat of Geltrex (10mg/mL, Gibco, A14133-02) on top, incubated in low oxygen incubators (5% CO2, 5% 02) and passaged every 3 to 4 days as single cells using Accutase (Stemcell, 07920).

### Generation of stable genetically modified and transfection of hESCs

Human ESCs were transfected with Lifeact-GFP, H2B-iRFP and H2B-mNeonGreen. To generate the Lifeact-GFP-Blast plasmid, GFP-coding sequence was amplified from Addgene plasmid #240804 with the coding sequence for Lifeact (MGVADLIKKFESISKEE) contained in the forward primer and inserted into the vector plasmid (Addgene #240803) digested with XhoI (Promega) and NotI (Promega). The mT2TP transposase and H2B-iRFP-Blast plasmids were generated as previously described^29^. H2B-mNeonGreen was obtained from Addgene (H2B-mNeonGreen-IRESpuro2 was a gift from Daniel Gerlich (Addgene plasmid #183745). To establish the Lifeact-H2B ESCs, 5 µg of mT2TP transposase plasmid was co-transfected with 5 µg of Lifeact-mScarlet-Puro plasmid and 5 µg of H2B-iRFP-Blast plasmid. Similarly, for introduction of H2B-mNeonGreen, 5 µg of mT2TP transposase plasmid was co-transfected with 5 µg of H2B-mNeonGreen-Blast plasmid. 500.000 hESCs were passaged as single cells, washed twice with OptiMEM (Gibco, 12559099), and transfected using the NEPA21 electroporator (Nepa Gene). Cells were immediately transferred to PXGL medium with 10 µM Y27632 and incubated on a 6-well plate with MEFs and Geltrex. 2 days later, cells were either selected using Puromycin (2 µg/mL; Invivogen), Blasticidin (100 µg/mL; Invivogen) or sorted using FACS to create polyclonal lines. Lines were checked for their fluorescent reporter expression every week by checking their fluorescence under the fluorescent microscope. Lines were eventually FACS sorted on their fluorescence expression to exclude negative cells that have silenced the construct.

### Blastoid generation and expanded culture

hESCs and underlying MEFs were dissociated by incubation with Accutase (Stemcell, 07920) for 5 minutes at 37 °C followed and mechanical separation to create single cells. To exclude MEFs, one well of dissociated cells was cultured on a gelatin-coated, equally sized well for 60 minutes in PXGL medium. Hereafter, supernatant containing the stem cells was removed, spun down and put in N2B27 basal medium containing 10 µM Y27632 (Stemcell, 72304) and counted. The cells were seeded in Aggrewell microwell plates (Stemcell, 34415). Per well, 54.000 cells were seeded into 500 µL N2B27 basal medium with 10µM Y27632 and incubated for 24 hours at 5% O2. Then, medium was replaced by 2 mL N2B27 PALLY medium containing PD0325901(1 µM), A83-01 (1 µM, MedChemExpress, HY-10432), 1-oleoyl lysophosphatidic acid sodium salt (LPA) (5 µM, Tocris, 3854), hLIF (10 ng ml−1) and Y-27632 (10 µM). 48 hours later, the medium was replaced with N2B27 + LPA and after another 48 hours the blastoids were formed. For expanded culture without endometrium, blastoids were put on a 9.4 cm non-adherent petri dish (Greiner, 632180) in 4 mL IVC medium, consisting of adDMEM/F12 Life Technologies), supplemented with 1 × GlutaMAX (11574466, ThermoFisher Scientific), 20% FBS (A5256701, ThermoFisher Scientific), 1X ITS-X (Gibco, 10524233), NAC (Sigma-Aldrich, A9165-5G), 1x HEPES (11560496,

ThermoFisher Scientific), 1x non-essential amino acids (11140050, Gibco) , 100 U/mL Penicillin–Streptomycin (11548876, ThermoFisher Scientific) and 0,22% sodium lactate (041529.AK, ThermoFisher Scientific). Blastoids were cultured for an additional 2 days in the incubator on a shaker (20 rpm) to prevent them from sticking to the bottom or to each other. Then, blastoids were used for imaging or gene expression analysis.

### Co-culture of endometrium and blastoids

Blastoids and endometrium were co-cultured in misc. medium. To prevent an osmotic shock, blastoids were slowly acclimatized to FBS by increasing the concentration by 1% per 10 minutes before adding them to the co-culture. Around 5-10 blastoids were cultured within one well of an 8-well ibitreat high dish to prevent aggregation and nutrient depletion of the culture medium. Co-cultures were either used for live imaging in a controlled environment (5% CO2, 20% O2, 37 °C) or in an incubator with the same environment for 2-3 days. For the extended co-culture of 8 days, IVC medium was refreshed every 2 days.

### qPCR analyses

RNA was isolated using the RNAEasy kit (Qiagen) following the manufacturer’s instructions. Reverse transcription was performed on 250-500 ng RNA using the High-Capacity RNA-to-cDNA kit (ThermoFisher). qPCR analysis was performed using SYBR green (Bio-Rad) on a CFX384 Touch Real-Time PCR detection system (Bio-Rad). qPCR data were analyzed using excel and plotted in GraphPad Prism.

### Histology

Organoids were dissociated from the Basement Membrane Extract (BME) by washing with 10 mL of ice-cold AdDMEM+++ per well, followed by centrifugation at 500×g for 5 min. Pelleted organoids were fixed in 10% formalin for at least 30 min, then processed for paraffin embedding through serial incubations in 70%, 96%, and 100% ethanol, xylene, and liquid paraffin. Sections of 4 µm thickness were cut, rehydrated, and subjected to hematoxylin and eosin (H&E), periodic acid–Schiff (PAS), or immunohistochemical staining. For immunohistochemistry, antigen retrieval was performed according to the antibody manufacturer’s instructions, followed by blocking in 1% bovine serum albumin (BSA; MP Biomedicals, 160069) in PBS. Primary antibodies included anti-ACTUB (sc-23950, Santa Cruz), anti-PAEP (HPA020108, Sigma Aldrich), anti-MFSD2A (PA5-21049, ThermoFisher Scientific) and anti-ASCT2 (ab237704, Abcam) applied overnight at 4 °C. After three PBS washes, sections were incubated with secondary antibody (rabbit anti-goat; Southern Biotech, 6160-01) for 1 h at room temperature, followed by three additional PBS washes.

Signal detection was performed using BrightVision poly-HRP anti-rabbit (Agilent, K400311-2) or poly-HRP anti-mouse (Agilent, K400111-2) and developed with 3,3′-diaminobenzidine (DAB) for 10 min. Finally, sections were dehydrated and mounted using Pertex®. Images were acquired with a DM4000 optical microscope (Leica) and processed using Olyvia software.

### Immunofluorescence

Fixation of the 2D endometrium and the blastoids was performed by adding 4% PFA (paraformaldehyde, Roth, 4979.1) for 15 minutes, followed by 1 hour incubation with 0,1%Triton X-100 (ThermoFisher, HFH10) + 5% BSA (Sigma-Aldrich, 05470-5G) in PBS on RT. Subsequently, primary antibodies were added in a solution of 0,01%Triton X-100 + 2% BSA in PBS, and the samples were incubated overnight at 4°C. The next day, secondary antibodies were added and incubated for 2 hours at room temperature. Then, samples were washed with PBS and kept at 4°C until imaging. Samples were imaged using the Leica Stellaris 8 White Light laser system, either with a 93x glycerol or a 20x dry objective.

### List of antibodies used

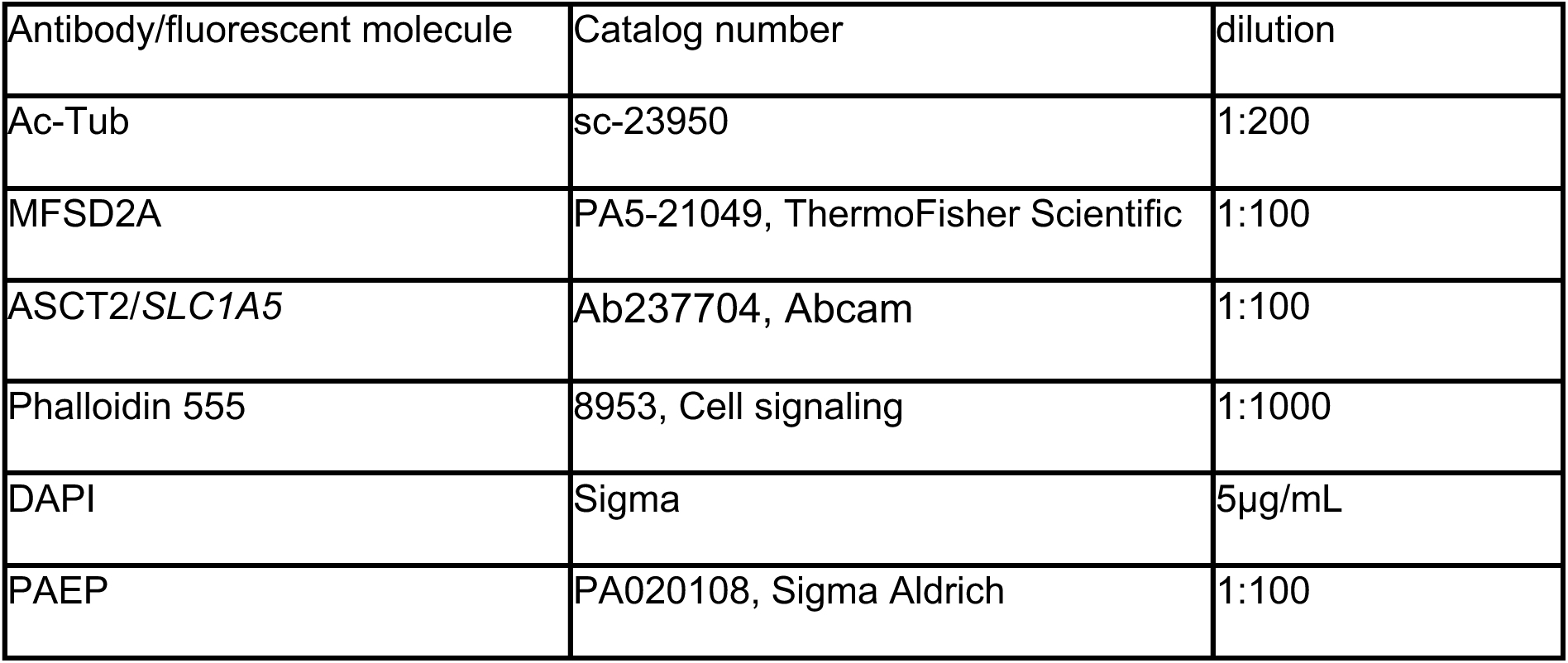

| Antibody/fluorescent molecule | Catalog number | dilution |
| --- | --- | --- |
| Ac-Tub | sc-23950 | 1:200 |
| MFSD2A | PA5-21049, ThermoFisher Scientific | 1:100 |
| ASCT2/SLC1A5 | Ab237704, Abcam | 1:100 |
| Phalloidin 555 | 8953, Cell signaling | 1:1000 |
| DAPI | Sigma | 5µg/mL |
| PAEP | PA020108, Sigma Aldrich | 1:100 |

### Live imaging

Live imaging of blastoids and endometrial co-culture was performed using a Leica Stellaris 8 White Light laser system. Imaging was performed in a humid atmosphere at 5% CO2, 20% O2 and 37°C. 488, 569 and 682 nm lasers were used for eGFP/mNeonGreen, m-Scarlet and iRFP, respectively, and images were either taken with a 93x glycerol or a 20x dry objective. For imaging, a field of view was chosen to focus on the interface of the endometrial epithelium and the blastoid. Z-Stacks of approximately 30 µm were taken with an increment of 1µm at 1024 x 1024 pixels every 30 minutes.

### Image data analyses

To measure the height of the endometrial epithelial cells, Z-stacks of differentiated 2D endometrial layers were taken with an increment of 1 µm. With the Fiji plugin orthogonal views, we generated a cross section. The distance from the basal to the apical side within individual cells was measured by drawing a line between the two sides in the middle of the cell. To quantify the H2B expression in blastoid and endometrial nuclei, the fluorescent intensity of H2B-iRFP and H2B-mNeon in nuclei from blastoids and endometrium were quantified by generating a maximum intensity projection, drawing a line through the cells covering the nucleus and measuring the intensity.

Surface area of attached blastoids was measured by manually defining a region of interest at the contact site of the endometrium and blastoids, surrounding the entire blastoid structure. Subsequently, the area was measured in Fiji.

### Expression of *SLC1A5* and *MFSD2A* in *in vivo* endometrium

To make the dot plots of *in vivo* fusion receptor expression, single cell RNAseq data from the non-pregnant uterus was downloaded from the reproductive cell atlas^25^ (https://www.reproductivecellatlas.org/non-pregnant-uterus.html). The dot plots were made using Scanpy’s (v1.11.4) built-in dotplot function grouping the cells by "Broad cell type" as annotated by the creators of the dataset.

### Bulk mRNA sequencing

RNA was isolated from 2D endometrial organoids, as done for qPCR analyses. Bulk mRNA sequencing was performed by Single Cell Discoveries (Utrecht, Netherlands). Briefly, polyA-enriched RNA was reverse transcribed and sequenced on an Illumina NextSeq500. Analysis of bulk RNA-seq samples was performed in R using DESeq2 (version 1.26.0)^31^.

### Statistics

Across all figures, error bars represent the standard error of the mean (SEM). Unpaired Student’s t-tests were used for significance testing. Statistical analyses and data visualization were performed using GraphPad Prism v10.6.0. The meaning of individual data points is detailed in the respective figure legends.

