## Extended Data for "Human embryo implantation involves Syncytin-2/MFSD2A-mediated heterokaryon formation with maternal endometrium"

### Extended data figures and legends

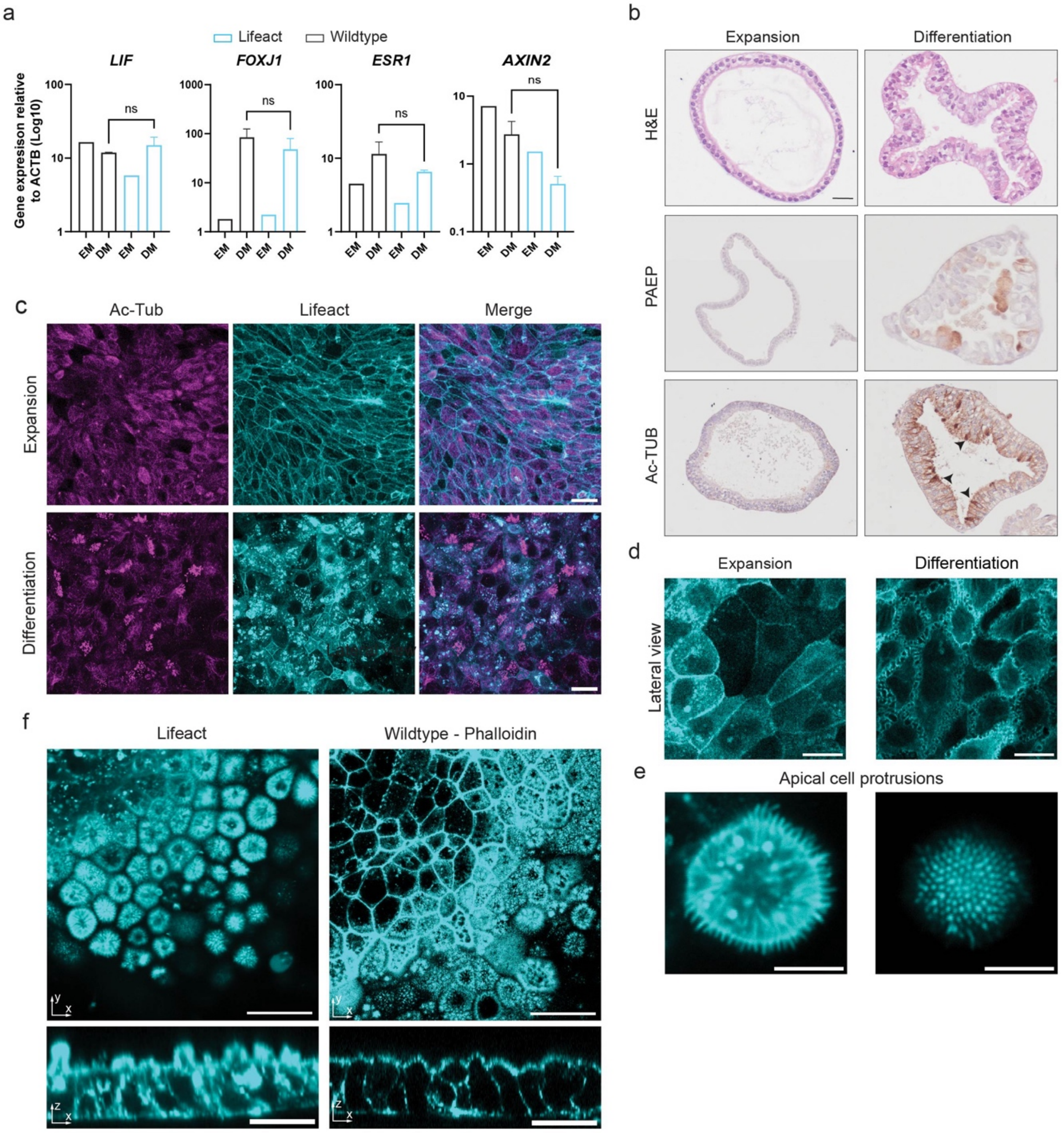

**Extended data figure 1- 2D endometrial epithelium expressing Lifeact-mScarlet differentiates upon stimulation with hormones.**

(**a**) Relative expression of endometrial marker genes measured by RT-qPCR: *LIF* (secretory cells), *FOXJ1* (ciliated cells), *ESR1* (hormone-responsive gene), *AXIN2* (Wnt target gene). Comparison between Wildtype and Lifeact-mScarlet organoids cultured in Expansion Medium (EM) or Differentiation Medium (DM). Data represent mean ± s.e.m.; statistical analysis by unpaired *t*-test, two tailed, ns= non-significant (n = 3). **(b)** Hematoxylin and eosin (H&E) staining and immunohistochemistry for PAEP and Acetylated-Tubulin in Lifeact-mScarlet expressing endometrial organoids cultured in EM or DM. Arrows indicate ciliated cell regions. Scale bar, 20 µm. **(c)** Immunofluorescence staining of ciliated cells (Ac-Tub, magenta) and Lifeact-mScarlet (cyan) in endometrial organoids cultured in EM or DM. Scale bar, 50 µm. **(d)** View of lateral membrane morphology in 2D Lifeact-mScarlet endometrial organoids. Left panel is in EM and right panel is in DM. Scale bar, 10 µm. **(e)** Apical membrane protrusions in 2D Lifeact-mScarlet endometrial organoids cultured in DM. Left image is a lateral view, right image is the top of the protrusion. Scale bar, 5 µm. **(f)** Top (upper panels) and orthogonal (lower panels) views of 2D Lifeact-mScarlet and Wildtype organoids stained with Phalloidin showing epithelial organization. Scale bar, 20 µm.

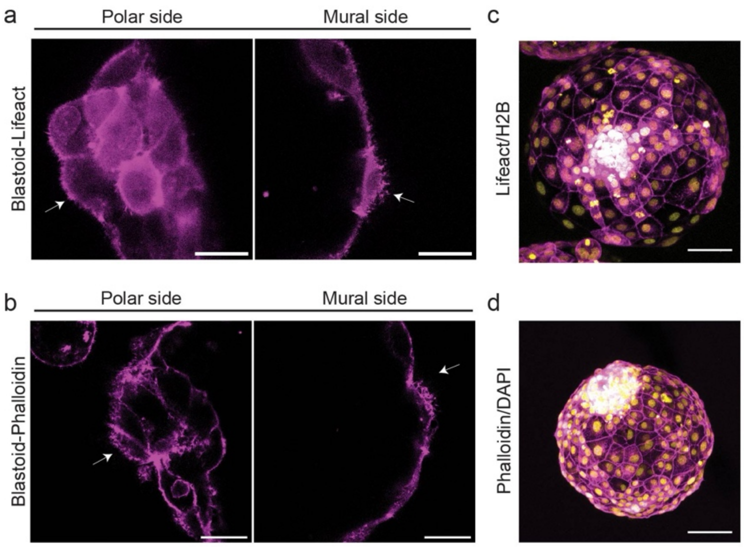
a
Polar side
Mural side
Blastoid-Lifeact
c
Lifeact/H2B
b
Polar side
Mural side
Blastoid-Phalloidin
d
Phalloidin/DAPI

**Extended data figure 2- Comparison of morphology between WT and Lifeact-GFP expressing blastoids**

**(a)** Images of Lifeact-GFP-expressing blastoids (magenta) viewed from the polar side (left) and mural side (right). Arrows indicate microvilli. Scale bar, 20 μm. **(b)** Images of Wildtype blastoids stained with Phalloidin (magenta), shown from the polar (left) and mural (right) sides. Arrows indicate microvilli. Scale bar, 20 μm. **(c)** Max-projection image of Lifeact-mScarlet (magenta) and H2B-iRFP (yellow) fluorescence in blastoids. Scale bar, 20 μm. **(d)** Max-projection image of Wildtype blastoid stained with Phalloidin (magenta) and DAPI (yellow). Scale bar, 20 μm.

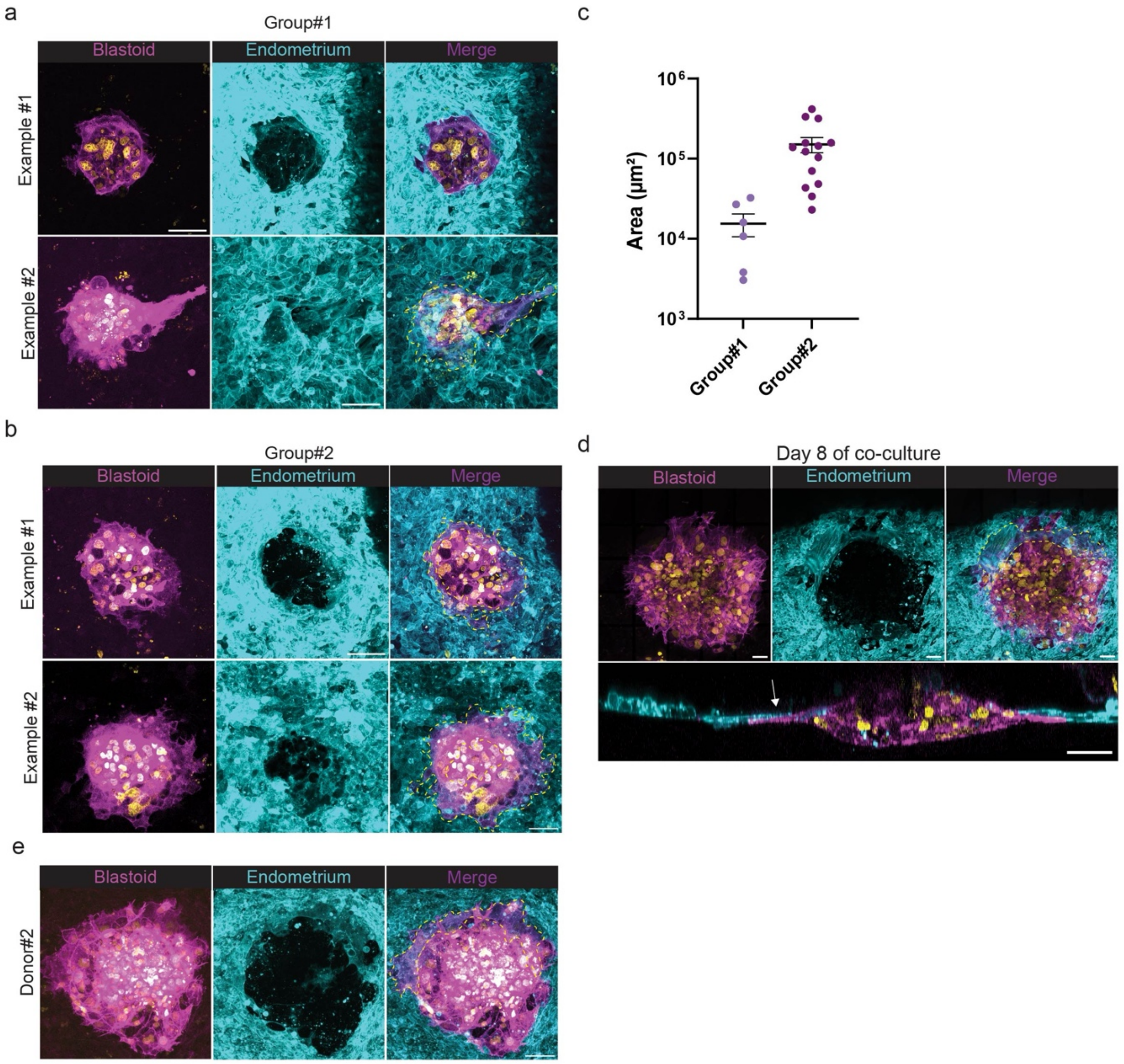
a
Group#1
Blastoid
Endometrium
Merge
Example #1
Example #2
c
10^6
10^5
10^4
10^3
Area (µm²)
Group#1
Group#2
b
Group#2
Blastoid
Endometrium
Merge
Example #1
Example #2
d
Day 8 of co-culture
Blastoid
Endometrium
Merge
e
Blastoid
Endometrium
Merge
Donor#2

**Extended data figure 3- Different groups of attached blastoids and confirmation of fusion with endometrium from additional donor.**

**(a)** Representative images showing two examples of early invasion during blastoid-endometrial fusion. From left to right: blastoid (magenta), endometrium (cyan), and merged image highlighting fused cells (circled in yellow). Max-projection of 3x1 µm planes. Scale bar, 50 µm. **(b)** Representative images showing two examples of group 2 of invasion during blastoid-endometrial fusion. From left to right: blastoid (magenta), endometrium (cyan), and merged image highlighting fused cells (circled in yellow). Max-projection of 3x1 µm planes. Scale bar, 50 µm. **(c)** Dot plot of the attached blastoid area ($\mu m^2$) between Group 1 and Group 2. Data represent mean ± s.e.m. **(d)** Representative images of the co-culture at day 8 showing fusion between the endometrium and blastoid. Top: top-view image of an attached blastoid with one multinucleated cell positive for Lifeact-mScarlet (cyan), Lifeact-GFP (magenta), and H2B-iRFP (yellow). Bottom: Orthogonal view of the same region with a white arrow indicating the giant fused cell. Max-projection of 5x1 µm planes. Scale bar, 50 µm. **(e)** Blastoid-endometrial fusion observed in an independent donor (Donor 2). From left to right: blastoid (magenta), endometrium (cyan), and merged image highlighting fused cells (circled in yellow). Max-projection of 3x1 µm planes. Scale bar, 50 µm.

a

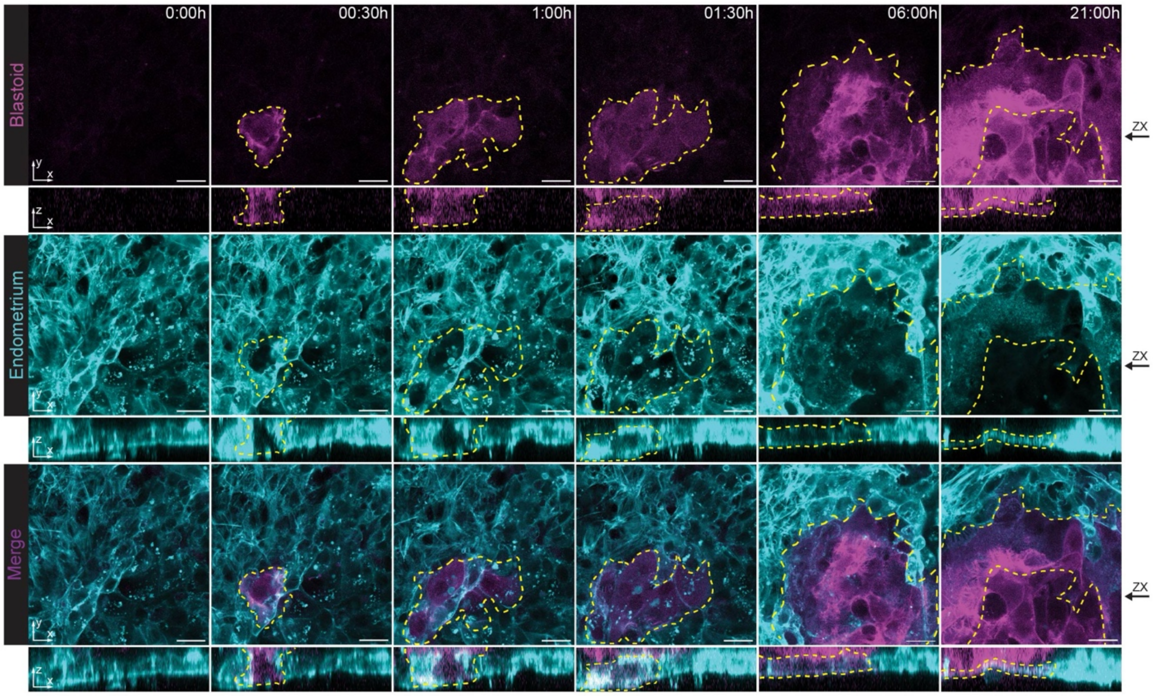
0:00h
00:30h
1:00h
01:30h
06:00h
21:00h
Blastoid
Endometrium
Merge
ZX

b

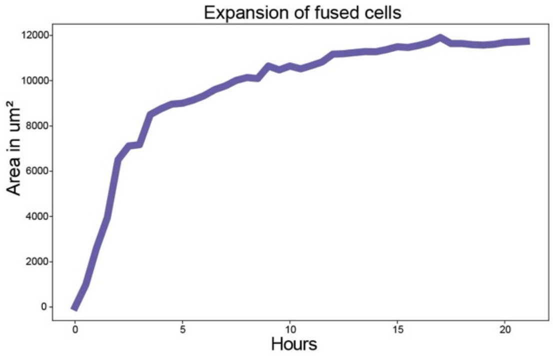
Expansion of fused cells
Area in um²
Hours
0
2000
4000
6000
8000
10000
12000
0
5
10
15
20

**Extended data figure 4- Extended live imaging images and quantification**

**(a)** Time-lapse imaging showing top and orthogonal views of blastoid-endometrial fusion. Top panels: blastoid (magenta); middle panels: endometrium (cyan); bottom panels: merged images highlighting fused cells (circled in yellow). Images represent a 1 µm z-plane. ZX represents the position of which the orthogonal view is generated. Scale bar, 20 µm. **(b)** Quantification of the fused cell area (µm²) of extended data figure 4a over time (in hours).

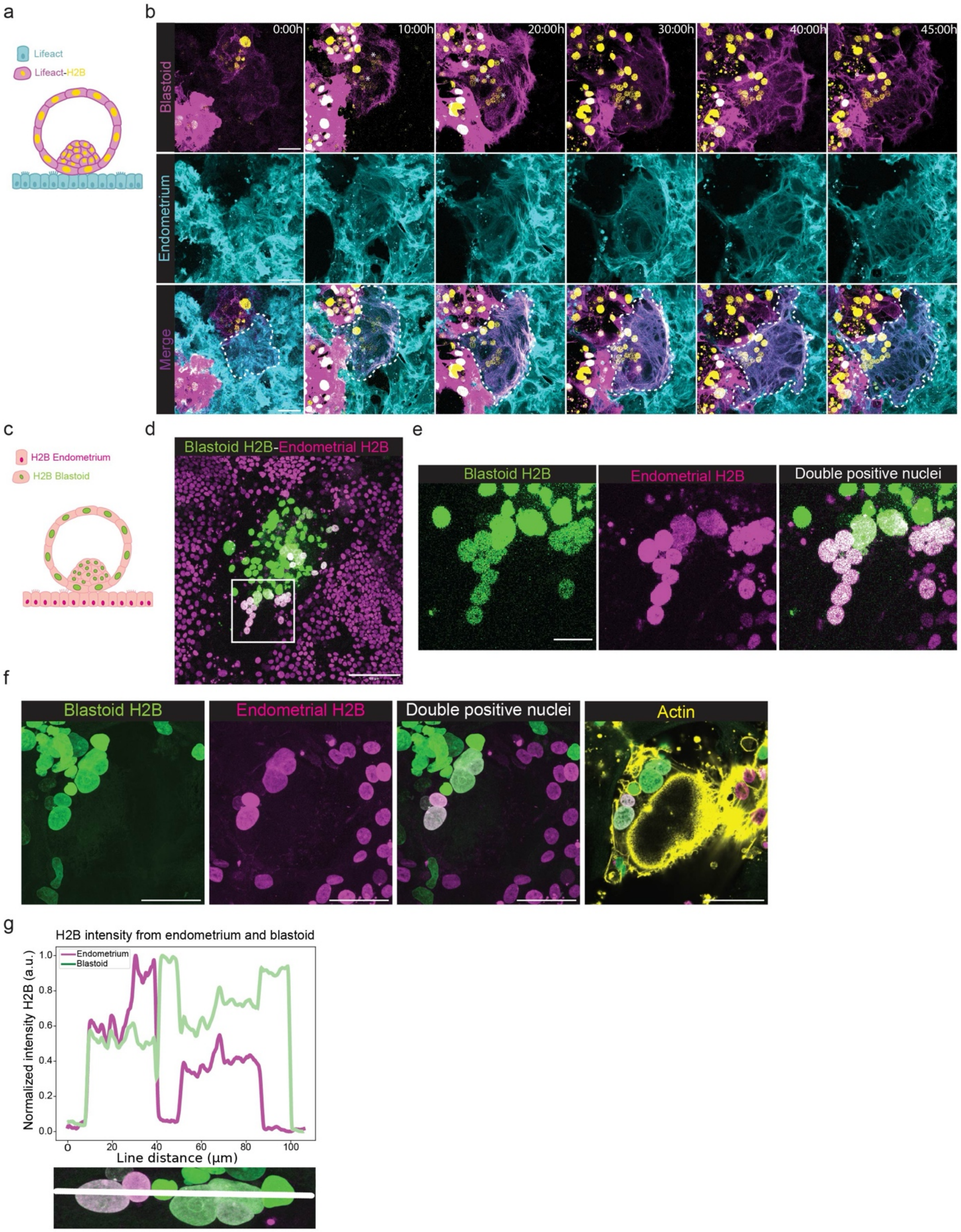

a
Lifeact
Lifeact-H2B
b
0:00h
10:00h
20:00h
30:00h
40:00h
45:00h
Blastoid
Endometrium
Merge
c
H2B Endometrium
H2B Blastoid
d
Blastoid H2B-Endometrial H2B
e
Blastoid H2B
Endometrial H2B
Double positive nuclei
f
Blastoid H2B
Endometrial H2B
Double positive nuclei
Actin
g
H2B intensity from endometrium and blastoid
Endometrium
Blastoid
Normalized intensity H2B (a.u.)
Line distance (µm)

**Extended data figure 5- Blastoids and endometrium nuclei share the same cytoplasm. Analysis and quantification.**

**(a**) Schematic of co-culture setup with Lifeact-mScarlet (cyan) endometrium, and Lifeact-GFP (magenta) and H2B-iRFP (yellow) blastoids. **(b)** Time-lapse imaging of blastoid-endometrial fusion. Top panels: blastoids (magenta); middle panels: endometrium (cyan); bottom panels: merged images showing fused cells (circled in white). Scale bar, 50 μm. **(c)** Schematic of co-culture setup with H2B-iRFP (magenta) endometrium and H2B-mNeon (green) blastoids. **(d)** Merged image showing nuclei from H2B-iRFP (magenta) endometrium and H2B-GFP (green) blastoids. Scale bar, 0 μm. **(e)** Zoom-in of d (white box). From left to right: H2B-GFP (green) blastoids, H2B-iRFP (magenta) endometrium, and merged image showing double-positive nuclei (white). Scale bar, 20 μm. **(f)** Images showing (from left to right): H2B-GFP (green) blastoids, H2B-iRFP (magenta) endometrium, merged image with double-positive nuclei (white), and merged image with actin staining (yellow). Scale bar, 50 μm. **(g)** Line-scan analysis of H2B fluorescence intensity in the endometrium (magenta) and blastoids (green). X-axis: distance (μm); Y-axis: normalized intensity (a.u.). The image below represents the region used for intensity measurement.

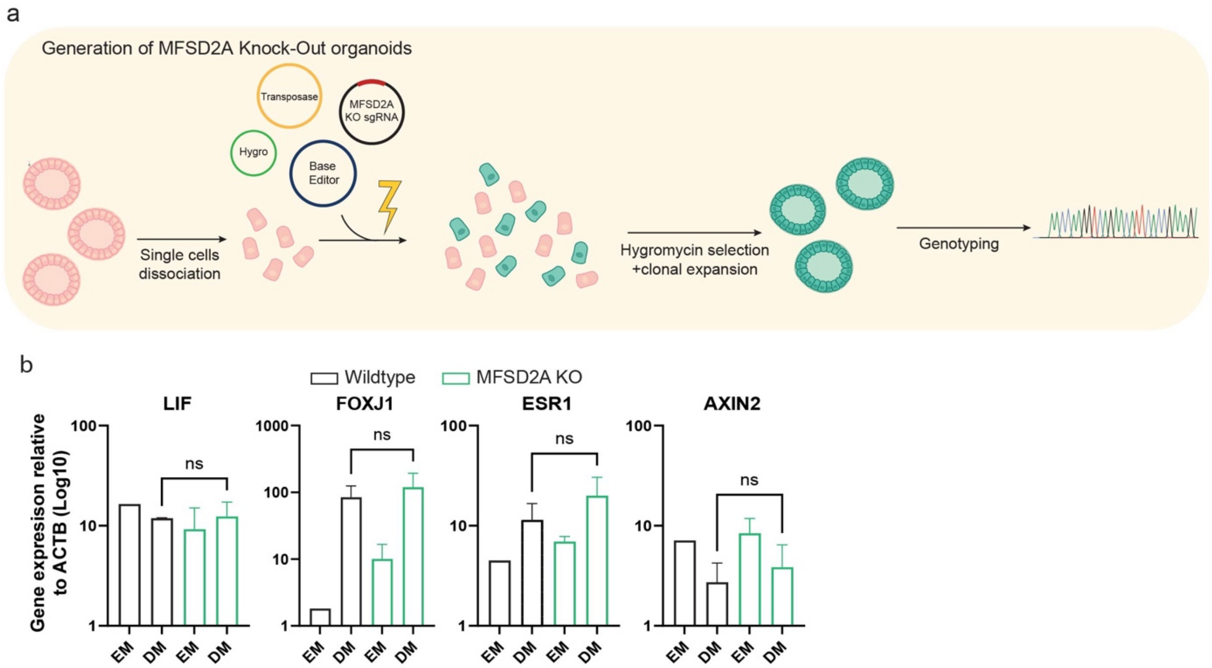
a
Generation of MFSD2A Knock-Out organoids
Transposase
MFSD2A KO sgRNA
Hygro
Base Editor
Single cells dissociation
Hygromycin selection +clonal expansion
Genotyping
b
Wildtype
MFSD2A KO
LIF
FOXJ1
ESR1
AXIN2
Gene expression relative to ACTB (Log10)
ns
EM
DM

**Extended data figure 6-Generation and validation of MFSD2A knockout endometrial organoids.**

**(a)**Schematic of the generation of MFSD2A KO lines in endometrial organoids using CRISPR-BaseEditors. **(b)** Relative expression of endometrial marker genes measured by RT-qPCR: *LIF* (secretory cells), *FOXJ1* (ciliated cells), *ESR1* (hormone-responsive gene), *AXIN2* (Wnt target gene). Comparison between Wildtype and MFSD2A KO organoids cultured in EM or DM. Data represent mean ± s.e.m.; statistical analysis by unpaired *t*-test, two tailed, ns= non-significant (n = 3).
